# Transgenic Mice and Pluripotent Stem Cells Express EGFP under the Control of miR-302 Promoter

**DOI:** 10.1101/450791

**Authors:** Karim Rahimi, Sara Parsa, Mehrnoush Nikzaban, Seyed Javad Mowla, Fardin Fathi

**Affiliations:** Department of Molecular Biology and Genetics, Aarhus University, Aarhus, Denmark; Molecular Genetics Department, Faculty of Biological Sciences, Tarbiat Modares University, Tehran, Iran; Department of Biological Sciences and Biotechnology, Faculty of Science, University of Kurdistan, Sanandaj, Iran; Cellular and Molecular Research Center, Research Institute for Health Development, Kurdistan University of Medical Sciences, Sanandaj, Iran

**Keywords:** Embryonic Stem Cells, Transgenic Mice, *Mir-302* Promoter, Blastocysts, Somatic Tissues

## Abstract

MicroRNAs are a group of short non-coding RNAs that undertake various roles in different cell signaling pathways and developmental stages. They regulate gene expression levels at the post-transcriptional stage, which results in cleavage of mRNAs or repression of their translation. Some miRNAs, including the *miR-302* cluster, are critical regulators for the stemness state of embryonic stem cells and cell fate patterning. The *miR-302* cluster is located in the intron of a non-coding gene that has no other reported function, other than hosting *miR-302*, and grant a complex expression regulation through upstream its regulatory sequences. To date, analysis of the *miR-302* expression pattern in a transgenic mouse model has not been reported. In this study, we generated transgenic mice that expressed EGFP driven by *miR-302* upstream regulatory sequences that harbored the core promoter of its host gene. We examined the activity of the *miR-302* promotor in somatic tissues of transgenic mice, transgenic blastocysts, and embryonic stem cells derived from transgenic blastocysts. Our results showed that *miR-302* highly expressed in both blastocysts and the first passages of transgenic embryonic stem cells, and has low expression in the somatic tissues of transgenic mice. It could be concluded that different temporal and spatial gene expression patterns occur during the embryonic and adult stages in mice.

## Introduction

### MicroRNAs and their role in stemness and development

MicroRNAs (miRNAs) are short non-coding RNAs that execute post-transcriptional repression by directly annealing to the complementary sequences of their target RNA (Lewis et al., 2005) and participate in gene regulation (Yu et al., 2006). The functions of only a small group of miRNAs have been analyzed. The miRNAs participate in the regulation of different cell signaling pathways that include cell differentiation, proliferation, growth and development, and cell death, in addition to the immune system and cancer promotion or inhibition (Sayed & Abdellatif, 2011). They are considered to be vital for animal development, regulation of cell proliferation, and have an association with diseases (Wang, Stricker, Gou, & Liu, 2007).

In addition to transcription factors (TFs) (Finley et al., 2018; Niwa, 2018) and cell surface markers (Sahlberg et al., 2014), miRNAs are increasingly recognized as potential stem cell markers (Lou et al., 2018; Song et al., 2017). The miRNA expression, as with other genes, is under the control of TFs and their genes may be transcribed by RNA polymerase II or III. Studies have previously described their general genetic structure and the mechanism of their biogenesis and processing (Lee et al., 2003; Lee et al., 2004; Borchert et al., 2006; Winter et al., 2009; Olena & Patton, 2010).

The *miR-302a-d*/367 cluster (*miR-302s*) is transcribed from a Pol II promoter. The primary RNA consists of 2-3 exons in humans and 2 exons in mice. *MiR-302s* is a polycistronic miRNA cluster that includes *miR-302*b/c/a/d and *miR-367*. Their order within the cluster represents their placement within the gene. These miRNAs share the same promoter and are generated from the same primary transcript (Barroso-delJesus et al., 2008; Rahimi et al., 2018). All five miRNAs are produced from an intron sequence. *MiR-302a-d* are highly related and share the same seed sequence, which is also shared with *miR-290-295* in mice and *miR-373* in humans. Although *miR-367* is processed from the same intron sequence (Rosa & Brivanlou, 2009), and its seed sequence is not the same as the other miRNAs in the cluster and targets different sites in the transcriptome. *MiR-367* regulates *Smad7* and TGF-β signaling in human pancreatic cancer cells, which promotes invasion and metastasis of these cancer cells (Zhu et al., 2015). Also, *miR-367* induces proliferation and stem cell-like behavior in medulloblastoma cells (Kaid et al., 2015). *MiR-302* induce somatic cell reprogramming and inhibit the differentiation process in stem cells (Hu, Zhao, & Pei, 2013). Pluripotency factors Oct4, Sox2, and Nanog strongly regulate murine *miR-302* expression (Tian et al., 2011) and the expression of this miRNA cluster clearly follows the expression level of these TFs (Hu et al., 2013). The expression pattern and regulation of the *miR-302* gene is mediated by the Wnt/β-catenin signaling pathway (Bräutigam, Raggioli, & Winter, 2013). *MiR-302* functions by down regulation of the cell cycle regulators CDKN1A and RBL2 (Subramanyam et al., 2011), and the TGF-β signaling antagonist Lefty1/2 (Barroso-deljesus et al., 2011). Furthermore, *miR-302* regulates the expression of nuclear receptor subfamily 2, group F, member 2 (NR2F2) which suppresses Oct4 and Sox2 expressions (Rosa & Brivanlou, 2011). Overexpression of *miR-302* can promote somatic cell reprogramming (Anokye-Danso et al., 2011). The introduction of a combination of *miR-302*s with *miR-200c* and *miR-369s* can efficiently reprogram mouse and human somatic cells to the pluripotent stage, which demonstrates that *miR-302* not only plays a critical regulatory role in pluripotency maintenance but also contributes to somatic cell reprogramming (Miyoshi et al., 2011). It has been shown that ectopic expression of *miR-302* can be a powerful tool to mediate induced pluripotent stem cell (iPSC) reprogramming without the need for OCT4, SOX2, KLF4, and MYC TFs (Anokye-Danso et al., 2011).

During murine embryonic development, expression of the *miR-302* cluster is down regulated after day 8 of gestation as shown by complete embryo analysis (Rosa et al., 2009); however, little is known about its expression in individual poly- or pluripotent stem cells. RT-PCR and whole mount in situ hybridization indicate *miR-302* expression in the developing lungs of murine embryos up to day 15 of gestation. This expression, together with functional studies, suggests an active role for *miR-302* in proliferation and specification of the lung tissues (Tian et al., 2011). Adult stem cells are multipotent cells that differentiate to mature somatic cells of the tissues in which they are located. They reside in a particular microenvironment niche. MiRNAs are key regulators in the self-renewal and differentiation process of adult stem cells (Mathieu & Ruohola-Baker, 2013). In humans, the *miR-302* cluster is located in the first intron of its host gene on chromosome 4. Embryonic stem cell (ESC) specific expression of the *miR-302s* is conferred by its immediate upstream regulatory region, located within 525 bp upstream of the transcription start site (Barroso-delJesus et al., 2008; Barroso-delJesus et al., 2009).

Transgenic mice are reliable in vivo tools that can be used to investigate the important roles of different genes and pathways. If the gene of interest is expressed in different organs and tissues, its investigation is much easier in the transgenic model rather than an in vitro study. Identifying the function of the gene of interest in an individual tissue is the key advantage of investigating gene expression patterns in transgenic mice compared to in vitro studies (Lambert, 2016). The possibility of inserting any DNA sequence that contains the gene of interest into the germline genome makes the mouse a powerful tool in basic biological investigations. By utilizing this technique, the inserted oligo sequence could carry a mutation or any changes in the genome sequence that results in a customized change in expression (Bouabe & Okkenhaug, 2013; Kitazawa, Medeiros, & LaFerla, 2012). In order to analyze the expression pattern of a gene, its upstream regulatory sequences could drive the expression of a fluorescent marker including *EGFP*. This transgene reporter allows in vivo and in vitro monitoring of the gene of interest. Transgenic mice act as a valuable model for developmental studies in that they can be used to track the expression profile of a gene of interest.

The *miR-302* cluster is highly conserved between mice and humans. The known promoter sequence contains binding sites for OCT4, SOX2, GATA6 and Nanog (Lewis, Burge, & Bartel, 2005), as TFs expressed in physiological and adult stem cells (Whissell et al., 2014; Amini, Fathi, Mobalegi, Sofimajidpour, & Ghadimi, 2014). The function of *miR-302* in establishing and maintaining stem cells is also documented by the observation that forced expression of individual members of the *miR-302* cluster can result in the formation of iPSCs (Kelley & Lin, 2012).

This project aims to use the expressions of stem cell specific miRNA *miR-302* regulatory sequences to visualize and show the expression pattern of the host gene of this miRNA cluster in transgenic mice tissues. This strategy uses the expression of the upstream regulatory sequences of the *miR-302* gene to express *EGFP*, which would generate green fluorescent stem cells.

## Materials and Methods

### Generation of “mmiR302pEGFP” transgenic mice that express *EGFP* under the transcriptional control of *miR-302*

The “hmiR302pEGFP” vector was a gift from Dr. Syed Jawad Mowla (Genetics Group, Tarbiat Modares University, Tehran, Iran). We sequenced the human *miR-302* (*hmiR-302*) cloned promoter and part of *EGFP* to confirm the vector sequence. To make this vector, the CMV promoter of the pEGFP-N1 vector (GenBank Accession No. U55762) was replaced by a human *miR-302* upstream regulatory sequence from -1,155 to +34 (chr4:112,648,667-112,649,855; UCSC Genome Browser for hg19) which contains the first 34 bp of the human *miR-302* exon 1. The CMV promoter was removed from pEGFP-N1 by using AseI and HindIII. After blunt ending and religation, the vector was re-digested with BglII and HindIII, and the promoter fragment was inserted (Figure S1). This vector was linearized with ApaLI for cell transfections.

We created the “mmiR302pEGFP” vector by digesting the pEGFP-C1 vector (GenBank; Accession No. U55763) with AseI and NheI to remove the CMV promoter. Murine *miR-302* immediate upstream regulatory sequences from -595 to +45 (chr3:127,544,494-127,545,132 - UCSC Genome Browser for mm9) were amplified by Pfu polymerase and the following primers: forward – 5’ AAGAATATTAATGTTTCCTGGTTGCTTCTAAT and reverse – 5’ AAGAATATTAATAATCACTAAATCAGGCAACC, which included AseI site and NheI site at the 5’ ends of the forward and reverse primers respectively. The PCR product was digested with AseI and NheI, then inserted in the linearized vector (Figure S2). Sequencing analysis was used to confirm the sequence of the insert. The vector was linearized with ApaLI for ESC transfection and pronucleus injections to generate transgenic mice.

In order to make transgenic mice, we used a purified and linearized “mmiR302pEGFP” vector for pronuclear microinjection into the mouse zygotes. Two-month-old C57bl6 female mice were superovulated by injections of 5 IU of PMSG and hCG (Sigma) at 48 h intervals. Fertilized eggs were collected from the oviducts of females that had mated with C57bl6 males. The linearized DNA fragments were injected into the pronuclei, and the injected eggs were transferred to 0.5-day pseudopregnant NMRI mice. Forward: 5’ GTTTCCTGGTTGCTTCTAAT and reverse: 5’ GCTGAACTTGTGGCCGTTTA primers were used for transgenic mice genotyping. All animal experiments were conducted in accordance with the Guidelines for the Care and Use of Laboratory Animals by Kurdistan University of Medical Sciences.

### Mouse Embryonic Stem Cell (mESC) culture and Electroporation

CJ7 mESCs (Swiatek & Gridley, 1993) were grown at 37°C in ESC medium that consisted of DMEM (Invitrogen, 10829-018), 15% fetal calf serum (Invitrogen, 12662-029), and 1000 U/ml LIF (Invitrogen, PMC9484) on mitotically inactivated feeder cells in 10 cm dishes for 2-3 days after seeding. When necessary, we used 0.1% gelatin cell culture dishes (Sigma-Aldrich, G1393).

Electroporation of the CJ7 ESCs was performed by re-suspending 1×10^7^ cells in 780 μl of PBS (pH 7.4; Invitrogen) in which the DNA had been dissolved. The prepared solution was transferred to an electroporation cuvette (Bio-Rad #165-2088). Electroporation settings were 240 V and 500 μF, and a time constant of approximately 6-8 seconds was set as an optimum condition. Electroporated cells were washed in 10 ml ESC medium and seeded on 4×6 cm dishes that had been pre-seeded with feeder cells. For all transfections, 25 μg of the vector was linearized in a mix of 50 μl of the FastDigest buffer, 10 μl of the enzyme, 25 μg of vector DNA, and ddH_2_O up to 500 μl, and incubated at 37°C overnight, followed by precipitation with ethanol. We started the G418 selection of the transfected cells 24-36 h after electroporation. Selected colonies could be picked after 6-8 days. The G418 concentration was approximately 350 μg/ml (Sigma-Aldrich, G9516). Mouth pipettes were used to choose the colonies, and each colony was transferred to the well of a round bottom 96-well plate that contained 50 μl of 0.05% trypsin-EDTA (Invitrogen, 25300-062). The selected colonies were incubated for 10 min at 37°C. Trypsin was inactivated by the addition of 100-150 μl ESC medium and the cells were transferred to a gelatinized (0.1% gelatin, Sigma-Aldrich, G1393) 96-well plate coated with feeder cells. Circular plasmids were used for transient transfection.

### In vitro fertilization and derivation of mESCs from blastocysts

The mice were superovulated by injections of 7.5 IU PMSG followed by an injection of 7.5 IU hCG 48 h later. The sperm collected from the caudal epididymides of mature, two-month-old “mmiR302pEGFP” transgenic mice was allowed to capacitate for 1 to 1.5 h at 37°C and subsequently diluted in HTF to a final concentration of 0.7-1.3×10^6^ sperm/mL. The collected oocytes from two-month-old *mmiR-302* transgenic mice were incubated with the spermatozoa for 4 h and then washed to remove any excess spermatozoa. The oocytes were cultured overnight in separate dishes that contained one drop of KSOM. After insemination, the obtained 2pn embryos were cultured in KSOM under mineral oil at 37°C in an atmosphere of 5% CO_2_ and air for 3 days until they reached the morula stage.

We isolated the mESCs by removing the zona pellucida of the morula embryos with acidic Tyrode’s solution. The obtained mESCs were cultured in KSOM+3i for 2 days. Next, the zona-free blastocysts were cultured in 3i+LIF medium for 5 to 7 days. Subsequently, the inner cell mass (ICM) cells were disaggregated with 0.05% trypsin that contained 0.2% EDTA after which the cell masses were placed on mitomycin C inactivated mouse embryonic fibroblast (MEF) cells in a 24-well plate with 3i+LIF medium that consisted of ESC medium supplemented with 0.8 uM PD184352, 2 uM SU5402, and 3 uM CHIR99021 (iSTEM medium; Takara, Japan). After 2-3 days, we observed the presence of compact cell colonies that resembled mESCs colony morphology. The ESC-like colonies, which expressed EGFP (Figure 4C) were picked up, trypsinized, and allowed to proliferate.

### Immunohistochemistry (IHC) for EGFP

After the mice were sacrificed, their somatic tissues were excised and maintained in PBS on ice. The samples were cut into pieces no larger than 1/4 cm^3^ and a piece of the sample was frozen on dry ice for RNA analysis. The remainder of the samples were partially fixed in 4% PFA for at least 4 h, but not longer than overnight if used for cryosections. Material for the cryosections was washed for at least 6 h in PBS and then incubated in 30% sucrose (in H_2_O) until the sample sank. Tissue-Tek (O.C.T. compound, Sakura) was used to freeze and embed the samples on dry ice. The frozen samples were stored at -80°C in a tightly closed vessel to prevent dehydration. Cryosections (10 μm thickness) were used for microscopic analysis. Rabbit anti-GFP polyclonal antibody (Bioss, bs-0890R) was diluted 1:100 in PBS + 1% FCS, and used as a primary antibody for staining of the tissue and ESCs. Alexa Fluor^®^ 488, goat anti-rabbit IgG (H+L, ab150077) was diluted 1:800 dilution in PBS + 0.1% FCS, and used as the secondary antibody.

## Results

### Human *miR-302* host gene expression analysis

The *hmiR-302* promoter element that included 1,155 bp of its gene regulatory sequences with a number of known TF binding sites upstream of the *EGFP* CDS (“hmiR302pEGFP” vector) was used for transient and stable transfection to test the activity of the *hmiR-302* promoter in different cell lines.

NT2 human embryonal carcinoma cells (Andrews, 1984) and HT1080 human fibrosarcoma cells (Benedict et al., 1984), as human cell models, and P19 cells, as the mouse cell model, were transiently transfected with the “hmiR302pEGFP” vector. In the NT2 and P19 cells, the *hmiR-302* promoter resulted in robust EGFP expression; however, there were fewer cells compared to the expression obtained with the CAG promoter, which was used as the positive control (Figure 1). In the HT1080 cells, the *hmiR-302* promoter resulted in the expression of the EGFP reporter (not shown), although this expression was weaker compared to NT2 cells.

**Figure 1:**
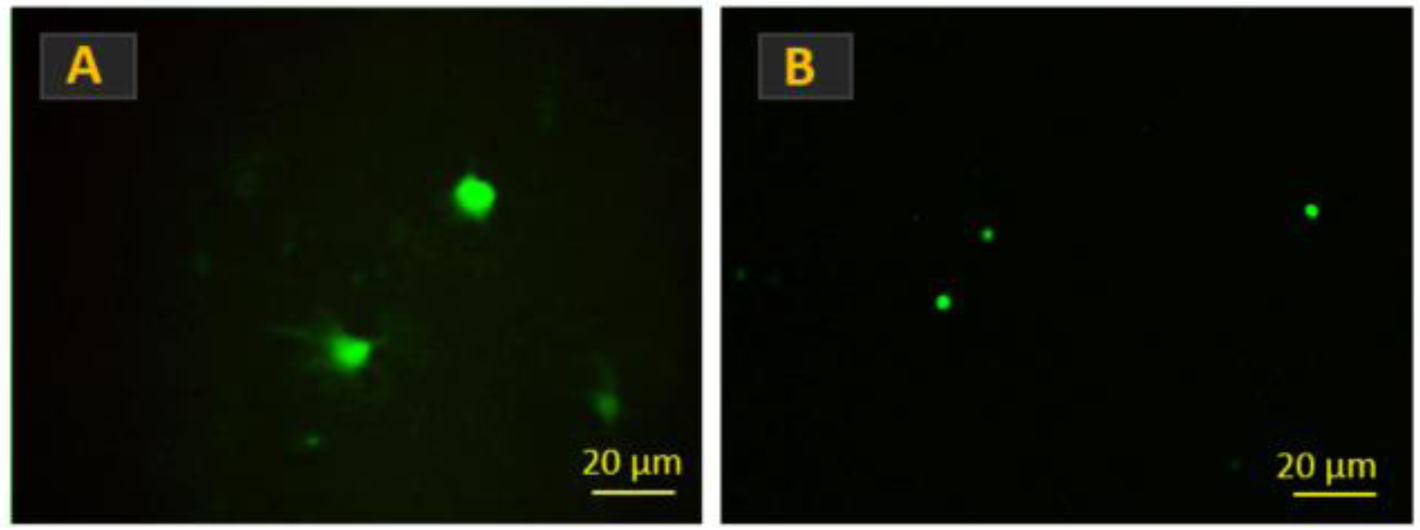
EGFP expression obtained from the transfection of NT2 and P19 cells driven by the *hmiR-302* promoter (hmiR302pEGFP). In NT2 cells, expression from the *hmiR-302* promoter (A) was comparable to the expression obtained from the CAG promoter (not shown), as the positive control. In P19 cells, the *hmiR-302* promoter only expressed in a few cells (B).

Stable transfected CJ7 mESCs cells (Swiatek & Gridley, 1993) with the “hmiR302pEGFP” vector were selected by using G418. The *hmiR-302* promoter induced visible EGFP expression in mESCs. Colonies with the best signals were picked and genotyped by transgene specific primers, and also tested by Southern blot using an *EGFP* probe that confirmed the genomic integration (data not shown). In order to test if the expression of the *hmiR-302* reporter depended on the pluripotent state of the cells, we removed the leukemia inhibitory factor (LIF) from the cell culture medium to induce differentiation. After removal of LIF, we noted that the EGFP signal turned off in the cells located at the margin of the colonies, which indicated that differentiation of the cells resulted in silencing of the *hmiR-302* promoter (Figure 2).

**Figure 2:**
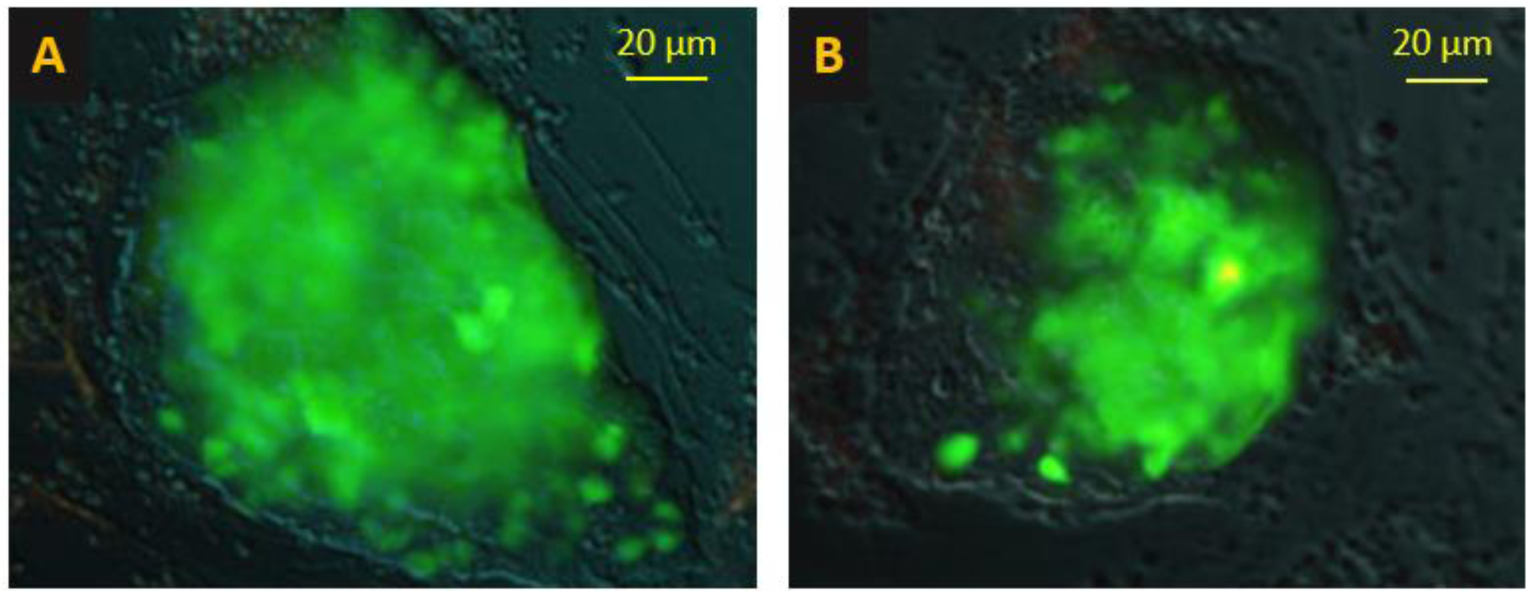
Stable transfection of mouse embryonic stem cells (mESCs) with the “hmiR302pEGFP” vector. The *hmiR-302* promoter induced clearly visible expression of EGFP in mESCs (A). We removed the leukemia inhibitory factor (LIF) from the medium and observed that the margin of the colonies began to differentiate, with reduced *miR-302* expression. Images were acquired 3 days after culture.

### Mouse *miR-302* regulatory sequences

In a separate study, we found the potential upstream regulatory sequences, the total structure of the gene cassette, and splice variants of the *mmiR-302* host gene (Rahimi et. el., 2018). In humans, the core promoter drives the gene and contains binding sites for some of the known stem cell specific TFs including OCT4, SOX2, and NANOG (Figures S3 & S4). In mice, the gene contains two exons, and the miRNAs are located in the intron. Since there is no open reading frame for the spliced transcript, it is anticipated that the gene is non-coding and only hosts the expression of the *miR-302* cluster. This gene overlaps the *Larp7* gene in the complimentary DNA strand.

### Transfection of CJ7 mESCs using the “mmiR302pEGFP” vector

The linearized “mmiR302pEGFP” plasmid was used for electroporation of CJ7 cells. Transfected cells expressed EGFP, which indicated the activity of the *mmiR-302* core promoter that drives the *EGFP* cassette (Figure 3A). After selection of the colonies with the greatest amount of green fluorescence, we performed transgene specific PCR genotyping and Southern blot analysis with an *EGFP* specific probe, which confirmed the stable integration (not shown). In a duplicate culture, the stably transfected cells were grown in differentiation medium, which resulted in down regulation of EGFP expression (Figure 3B).

**Figure 3:**
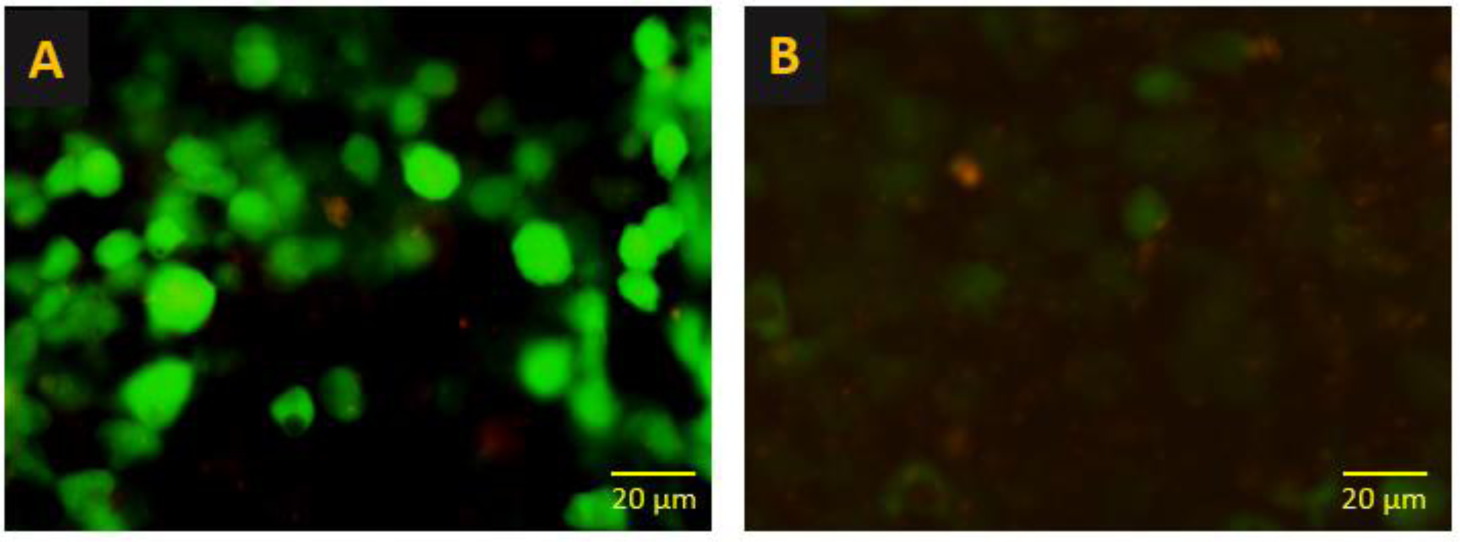
Stable transfection of mouse embryonic stem cells (mESCs) with “mmiR302pEGFP” vector. The 595 bp upstream genomic region of *mmiR-302* drives expression of *EGFP* in CJ7 cells (A). Removal of leukemia inhibitory factor (LIF) shows a decrease in *mmiR-302* reporter expression in the cells. Images were acquired 5 days after seeding.

**Figure 4:**
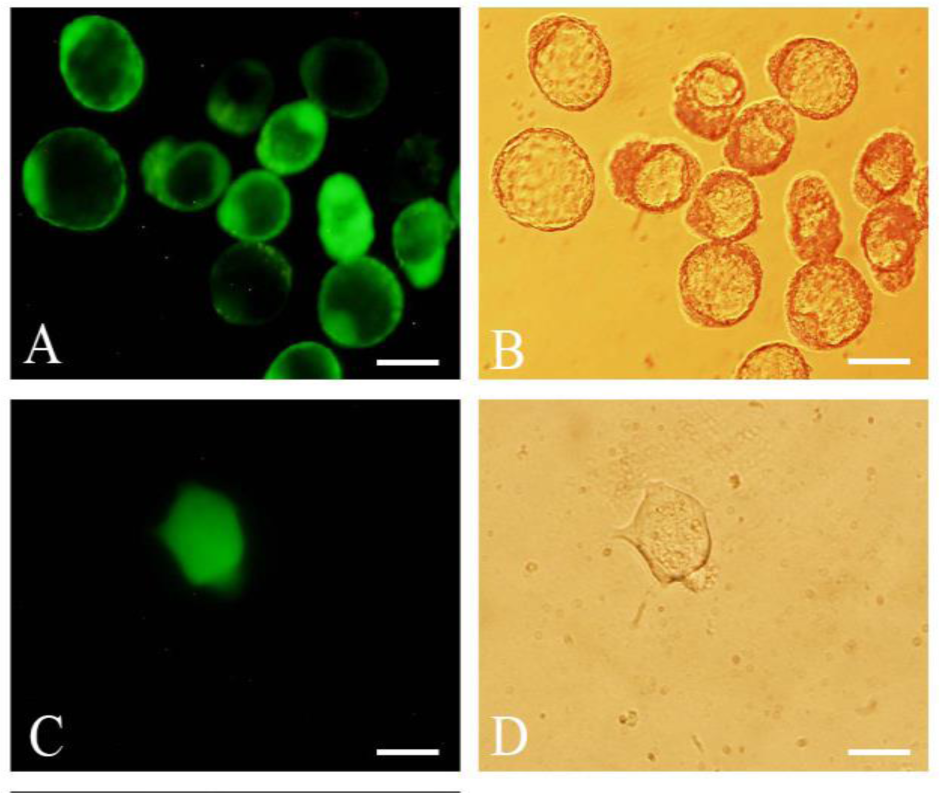
EGFP expression in the blastocysts obtained from transgenic mice (A). Isolated embryonic stem cells (ESCs) derived from blastocysts of transgenic mice expressed EGFP (C). B and D show the phase contrast counterparts of A and C, respectively. Scale bars: 50 μm.

### Generation of transgenic mice using the “mmiR302pEGFP” vector

Because the mESCs expressed the “mmiR-302pEGFP” reporter, we decided to make transgenic mice that utilized the same construct as for the *miR-302* expression study in mESCs. Transgenic mice were identified by PCR amplification of the transgenic cassette. RT-PCR analysis showed the expression of *EGFP* in some selected tissues derived from the transgenic mice (not shown).

### EGFP expression in the transgenic blastocysts and obtained mESCs

In order to test *mmiR-302* reporter expression in early embryogenesis, we analyzed the EGFP expression pattern in the transgenic morula, blastocysts, and blastocysts derived mESCs where *EGFP* was expressed by the *mmiR-302* core promoter. Transgenic morula and blastocysts, which were derived from IVF of transgenic oocytes and spermatozoa, had strong EGFP expression. Zona free blastocysts seeded in the 3i+LIF medium for 5 to 7 days resulted in ICM derived mESCs. Both transgenic single cells and 3-day obtained colonies showed significant EGFP expression. The transgenic cell colonies resembled mESC colony morphology (Figure 4).

We tested for EGFP expression in the somatic tissues of the transgenic mice independent of its fluorescence and used EGFP specific antibodies on cryosections of some of the transgenic tissues including heart, kidney, and liver. All of the tissues showed weak but distinct signal (Figure 5).

**Figure 5:**
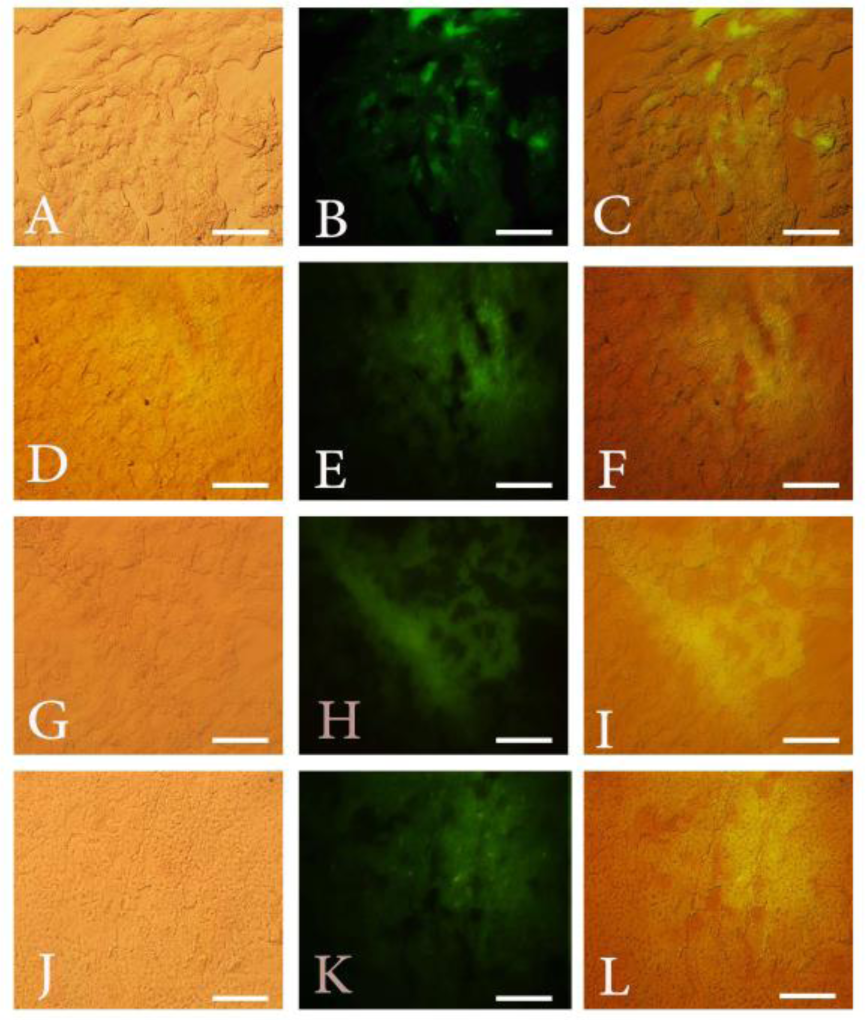
Immunostaining of some tissues in the “mmiR302pEGFP” transgenic mice. Heart tissue taken from an *EGFP* transgenic mouse was used as a positive control (A, B, and C). Heart tissue from “mmiR302pEGFP” transgenic mice (D, E, and F). Kidney tissue from “mmiR302pEGFP” transgenic mice (G, H, and I). Liver tissue derived from “mmiR302pEGFP” transgenic mice (J, K, and L). Scale bars: 100 μm.

## Discussion

Both stem cells and adult stem cells are known key factors for the renewal of different tissues. There are a number of stem cells markers. Some miRNAs regulate gene expression levels in the stem cells, such as the *miR-302* cluster. *MiR-302s* are located in the first intron of the host gene, which is classified as a non-coding RNA (Rahimi et al., 2018). The expression of this gene is driven by Pol II RNA polymerase, and its regulatory sequence is more complex than a simple promoter for miRNA expression. The expression complexity of the *miR-302* host gene indicates the involvement of this miRNA cluster in key cellular pathways and mechanisms. Since this gene does not code any protein, it was not possible to use an antibody. In this study, we used the *miR-302* host gene core promoter to express *EGFP* in the transgenic mice and tracked the expression pattern of this gene in different tissues. We could track the expression of *EGFP* in the blastocysts and isolated mESCs. We performed preliminary transient transfection experiments for both human NT2 and mouse pluripotent P19 and C17 stem cells to ensure the expression of the *miR-302* promotor and control reporter genes, and to answer the question of whether the promoter of *miR-302* was active in the mouse pluripotent EC and ESCs. P19 is a mouse teratocarcinoma cell line that has been derived from an ectopically transplanted early mouse embryo (McBurney & Rogers, 1982). This line represents a pluripotent stem cell that can differentiate in all three germ layers (Liu et al., 2011). The results of NT2 and P19 cells have agreed with the results of previous studies where the *miR-302* promoter was active in the cells used in the study (Hohjoh & Fukushima, 2007). *MiR-302* expression in the ESCs has been reported (Barroso-delJesus et al., 2008; Houbaviy, Murray, & Sharp, 2003).

Our results showed that the *miR-302* core promoter was active in the mESCs CJ7 line. We observed EGFP signals in undifferentiated ESCs cultured in medium with LIF. Removal of LIF from ESC growth medium led to the gradual disappearance of the EGFP signals. This could be indicative of the fact that *miR-302* plays a role in regulating the status of undifferentiated and pluripotent mESCs. Previous studies have reported the low expression of the *miR-302* gene in mESCs compared to mouse epiblast stem cells (Rosa & Brivanlou, 2009; Stadler et al., 2010). This disparity may be due to silencing of the transgene while randomly integrated into the genome or perhaps the *miR-302* expression has a different activity level in the different developmental stages. Of note, we considered making a better reporter by knocking-in the *EGFP* coding sequence in the last exon of the *miR-302* host gene in our study. However, the limiting factor was the generation of a knockout for the *Larp7* gene, a housekeeping gene located in the complement strand of the DNA where the *miR-302* host gene is placed.

The regulatory elements of human *miR-302* have already been reported (Barroso-delJesus et al., 2008). We began the expression study of the *miR-302* host gene by transfecting different ES and EC cell lines from humans and mice to analyze the expression level of EGFP driven by human *miR-302* host gene regulatory elements. Evaluation of mice and human *miR-302* promoter activity in CJ7 mESCs showed that the human promoter was more active.

In our study, transgenic mice were produced by injection of a plasmid vector that contained *EGFP*, which expressed under the activity of the mouse *miR-302* core promoter. Sperm and oocytes extracted from the transgenic mice were used for in vitro fertilization. ESCs were isolated from the resultant embryos that developed to the 4.5 day blastocyst stage. Significant EGFP signals in the embryo, particularly at day 5 of embryo development, indicated that the *miR-302* promoter was active in the embryo stage, which was consistent with the results of previous studies (Card et al., 2008). From the blastocysts that expressed the EGFP protein, we isolated 3 mESC lines that were expanded to the 10th passage. In the first and second passages, EGFP expression was observed in the isolated cells; however, expression gradually reduced so that it was not visible after the third passage, which could be the results of transgenic silencing due to the previously reported methylation (Mutskov & Felsenfeld, 2004). Immunostaining with an antibody against EGFP was performed in the passage-10 isolated ESCs (Figure S5). The positive results indicated less promoter activity in comparison to the initial passages and expression in the embryonic stage. Given the equal copy numbers of the *miR-302* promoter and EGFP protein in each green blastocyst and isolated mESCs, it was likely that the core promoter of *miR-302* was more active in embryos compared to embryo derived ESCs.

We examined *miR-302* promotor activity in various somatic tissues such as the liver, heart, kidneys, brain, bone marrow, and stomach that were extracted from “mmiR302pEGFP” transgenic mice. There were positive IHC results for EGFP in the liver, heart, and kidneys, which showed that the promoter was active in those tissues. In the brain, bone marrow, and stomach, the *EGFP* expression was only seen by RT-PCR (data not shown). Our results indicated low, but distinct *miR-302* promoter activity, in the somatic tissues while lack or low expression of *miR-302* has been reported in adult somatic cells and adult stem cells (Miyoshi et al., 2011; Wilson et al., 2009). Our result might be due to particular stem cells that resided in the tissues.

